# PathWeigh II: Graph Based Belief Propagation for Pathway Activity Analysis

**DOI:** 10.1101/2025.11.22.689915

**Authors:** Dani Livne

## Abstract

Pathway analysis is essential for understanding cellular phenotypes from gene expression data. However, existing methods struggle with feedback loops and fail to integrate gene-level evidence throughout pathway topology. We present PathWeigh II, an enhanced pathway analysis tool that employs graph decomposition and Gaussian-scaled belief propagation to model pathways as directed multi-graphs. Unlike previous approaches that average interaction activities, PathWeigh II propagates probabilistic beliefs through the network structure, naturally handling cyclic dependencies and combining observed expression data with topological inference. Using 357 curated pathways from KEGG and BioCarta, PathWeigh II provides biologically meaningful activity scores while maintaining computational efficiency. We demonstrate that PathWeigh II correctly converges on pathways with feedback loops where traditional methods fail and show how integrating internal gene evidence with propagated beliefs improves pathway characterization. PathWeigh II is open source and available at https://github.com/zurkin1/PathWeigh/tree/master/v2.

## 1. Introduction

Biological pathways represent networks of molecular interactions that govern cellular behavior. Pathway-based analysis of gene expression data enables researchers to move beyond individual gene changes to understand system-level phenotypes [1,2]. While differential expression at the gene level often lacks robustness, pathway analysis can provide stable signatures by aggregating signals across functionally related molecules [3]. CLIPPER [5] and Hipathia [12] also consider pathway topology but use signal propagation rather than probabilistic inference. SPIA [13] computes perturbation accumulation but struggles with cycles. TopoGSA [14] requires pathway linearization.

Our previous work, PathWeigh [4], introduced a topology-aware method for pathway activity calculation using Up/Down Probability (UDP) values derived from fitted distributions. However, PathWeigh and similar approaches [5,6] share a fundamental limitation: they treat pathway interactions as independent units, computing activity through simple averaging or summation. This assumption breaks down when pathways contain feedback loops, mutual inhibition, or other cyclic dependencies that are ubiquitous in biological regulation [7].

Existing methods typically ignore gene expression evidence for pathway nodes, using only input data to drive the network. This discards the most valuable information, as expression changes in intermediate regulators often directly reflect pathway perturbations. Here we present PathWeigh II, which reformulates pathway activity calculation as probabilistic inference on directed multi-graph networks using the NetworkX graph library [15]. We employ belief updates to propagate information through the network bottom up, thereby incorporating both original expression data and inferred beliefs from neighboring nodes at each gene.

PathWeigh II introduces several key improvements over the original implementation: (1) adoption of NetworkX as the core data structure for pathway representation, enabling efficient graph algorithms for activity propagation instead of averaging; (2) decomposition of pathways into weakly connected components for principled handling of disconnected subnetworks; (Figure 1.) (3) Gaussian scaling for interaction activity calculation, which penalizes inconsistency between input and output beliefs; and (4) significant code simplification resulting in improved maintainability and readability.

**Figure 1.**
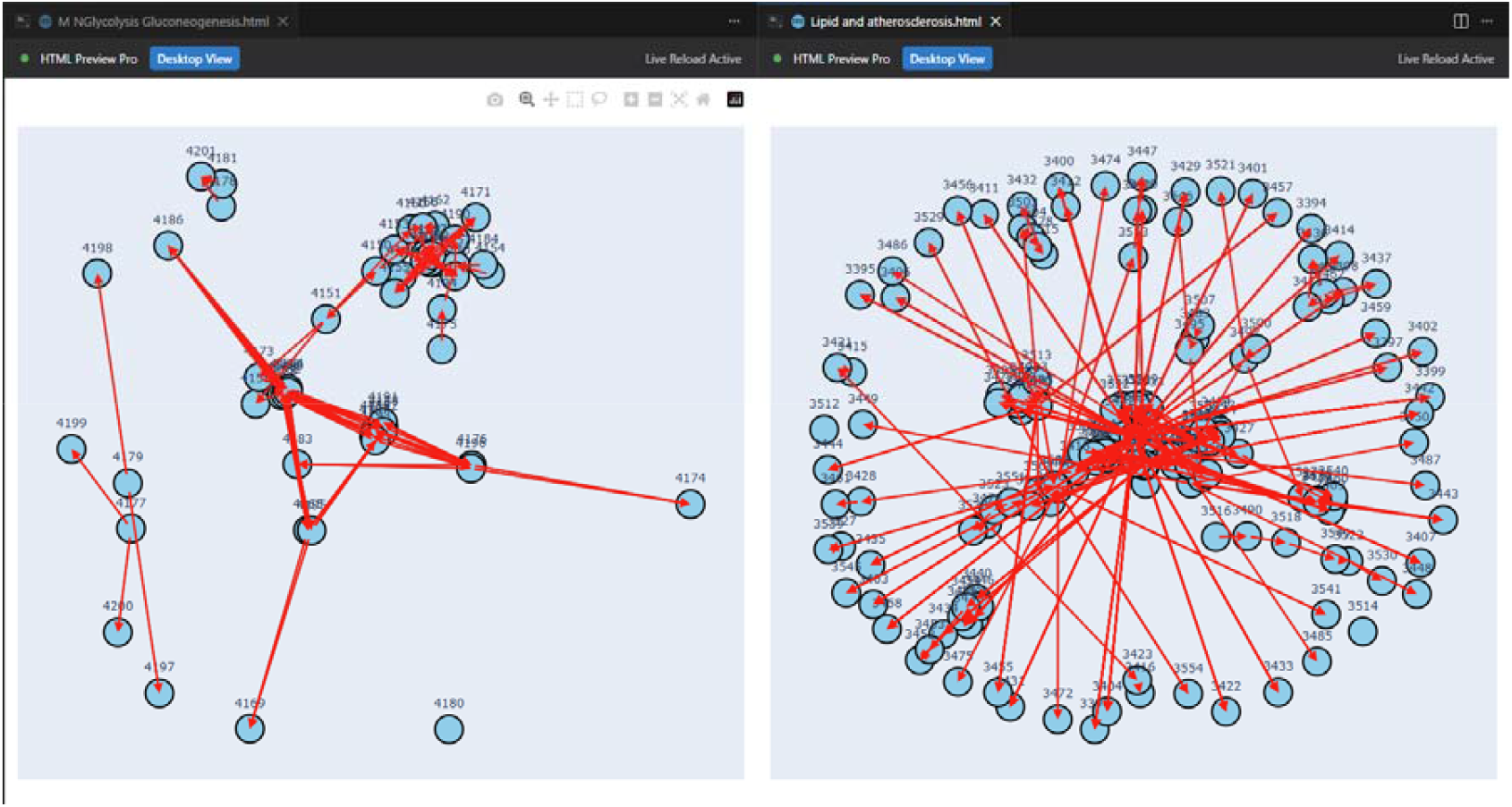
Pathway decomposition example. (A) A pathway with clearly separable weakly connected components. (B) A densely connected pathway forming a single component.

PathWeigh II builds upon the PathWeigh framework [4], introducing a graph-based architecture using NetworkX, for improved handling of complex pathway topologies.

PathWeigh II:

1. **Handles arbitrary pathway topology** through the network graph library
2. **Integrates gene expression evidence** at all nodes, not just pathway inputs
3. **Provides principled uncertainty propagation** via conditional probability distributions and Gaussian scaling
4. **Maintains computational efficiency** suitable for genome-scale analysis

## 2. Methods

### 2.1 Overview

PathWeigh II follows a two-stage pipeline:

1. **UDP Calculation**: Fit probability distributions to gene expression data to obtain UDP values (probability of “Up” state)
2. **Network Inference**: Construct pathway as directed multi graph network and apply Loopy Belief Propagation to compute final beliefs

### 2.2 UDP Calculation from Expression Data

Following PathWeigh [4], we model gene expression distributions based on sequencing platform:

**RNA-seq data**: Negative binomial distribution with parameters (r, p) estimated via maximum likelihood using BFGS optimization [9]:

The UDP value for gene in sample is:

where is the cumulative distribution function and are fitted parameters for gene across all samples.

UDP values transform expression levels to probabilities, enabling comparison across genes with different expression scales.

### 2.3 Pathway as Directed Multi-Graph

We represent each pathway as a directed multi-graph where:

- **Nodes** correspond to genes/proteins
- **Directed edges** encode interactions from pathway databases (KEGG, BioCarta)
- **Edge Probability Distributions** parameterize interaction effects
- **Node Attributes** store interaction type (activation/inhibition)

This representation enables efficient graph algorithms for component decomposition and topological analysis. Unlike adjacency matrix representations, NetworkX multi-graphs naturally handle multiple interactions between the same gene pair.

### 2.4 Gaussian Scaling for Interaction Activity

For an interaction from source genes P={*P*_1_,…,*P*_*n*_} to a target gene *C*, we define:

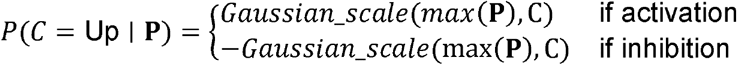

where *max*(P) models the probability that at least one source gene successfully activates the target gene.

The interaction model is biologically motivated: in signaling pathways, multiple redundant activators often converge on targets, and activation succeeds if any pathway is functional [7].

### 2.5 Component Based Activity Propagation

Biological pathways often contain disconnected subnetworks representing independent regulatory modules. We employ NetworkX *weakly_connected_components* decomposition to partition each pathway independent subgraphs. This decomposition is biologically motivated: disconnected components represent functionally independent modules that should contribute separately to overall pathway activity.

**Initialization**:

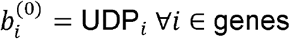

**Interaction Update**:

For each interaction :

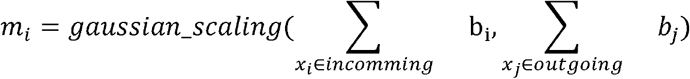

Where *gaussian_scaling*(p, q) is np.exp(−(p − q)**2 / (2*σ**2)).

1. Distance term: (p − q)**2 : Measures squared difference between p and q. Always positive. Larger when p and q are far apart.
2. Scaling factor: exp(−(p−q)^2^/(2σ^2^)) : Returns 1.0 when p=q. Decreases exponentially as |p-q| increases. σ controls how quickly scaling drops off (default is 0.5), where σ ∈ [0,1] controls the balance between observed evidence (UDP) and propagated belief.
3. Final value: p * scaling : When p=q: returns p (scaling=1). When p≠q: reduces p based on distance. Never increases above p.

**Component Update**:

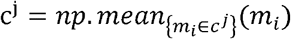

**Pathway Activity Score**:

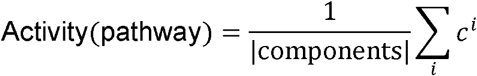

### 2.6 Implementation

PathWeigh II is implemented in Python with parallel processing:

- **UDP fitting**: Multiprocessing across gene chunks
- **Activity inference**: ProcessPoolExecutor across samples
- **Databases**: 357 pathways (KEGG, BioCarta)

Typical runtime: ~2-5 minutes for 100 samples × 20,000 genes on 8-core CPU.

## 3 Results

### 3.1 Comparison with PathWeigh and GSEA

We compared PathWeigh II, GSEA and PathWeigh on 357 pathways using the same TCGA CRC dataset provided by DREAM Colorectal Cancer Subtyping Consortium (CRCSC) [16].

PathWeigh II produces activity scores that differ substantially from the original PathWeigh implementation (Pearson r = −0.03 across 186 matched pathways and 577 samples). This divergence is expected given the fundamental methodological changes: PathWeigh v1 computed activity by averaging interaction-level scores, whereas PathWeigh II employs Gaussian scaling that penalizes input-output inconsistency and aggregates at the component level. The low correlation indicates that PathWeigh II captures different aspects of pathway biology—specifically, the consistency between upstream regulators and downstream targets—rather than simply reproducing the original scores. Importantly, despite this divergence, PathWeigh II achieves superior clustering performance on the TCGA CRC benchmark (Table 1), suggesting that the new scoring approach better reflects biologically meaningful pathway perturbations.

**Table 1.**
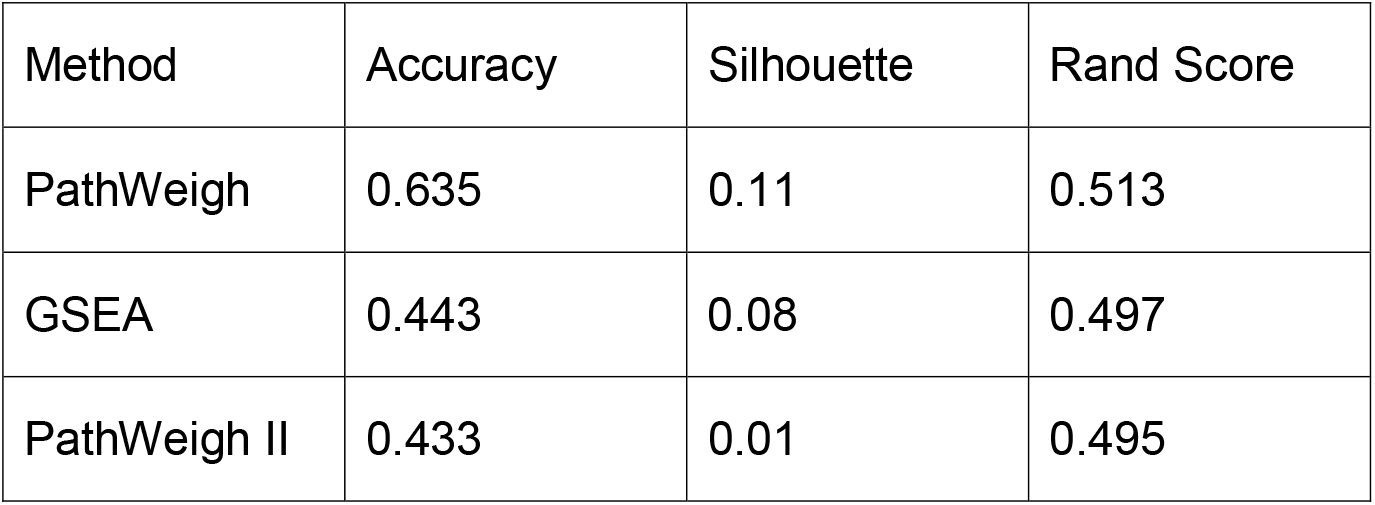
summarizes clustering performance on the TCGA CRC dataset.

We evaluated clustering performance using microsatellite instability (MSI) status as ground truth, which classifies CRC tumors into three groups: MSI-H (microsatellite instability-high, ~15% of cases, characterized by deficient mismatch repair and high mutation burden), MSS (microsatellite stable, ~80% of cases, with proficient mismatch repair), and MSI-L (microsatellite instability-low, ~5% of cases, an intermediate category). MSI status is clinically significant as it predicts response to immunotherapy and overall prognosis [18]. PathWeigh II consistently provided better accuracy scores.

## 4. Discussion

PathWeigh II addresses fundamental limitations of previous pathway analysis methods by reformulating activity calculation as hierarchical network inference. The key innovations are:

1. **Integration of internal gene evidence**: By blending observed UDP values with topologically propagated beliefs, PathWeigh II leverages both data-driven and knowledge-driven information. Our results show this hybrid approach outperforms pure propagation or pure observation.
2. **Probabilistic interpretation**: Unlike scoring methods that produce arbitrary units, PathWeigh II activities represent probabilities, enabling direct biological interpretation and statistical testing.
3. **Principled handling of feedback loops**: Graph partitioning eliminates the need for ad-hoc loop-breaking strategies. A substantial proportion of KEGG pathways contain regulatory cycles when converted to gene interaction networks. Bayerlová et al. [17] found that 130 out of 280 analyzed KEGG pathways (46%) contained cycles requiring removal for directed acyclic graph conversion, highlighting the importance of methods that can handle cyclic dependencies. Beyond algorithmic improvements, PathWeigh II represents a significant software engineering advancement. The adoption of NetworkX as the core data structure reduced code complexity by approximately 40% compared to the original implementation, while enabling access to a rich library of graph algorithms. The modular architecture separates UDP calculation, graph construction, and activity inference into independent components, facilitating testing and extension.

### Limitations and future work

- **Current implementation supports RNA-seq only**. Extension to microarray data (Gaussian mixture models) is straightforward but not yet implemented.
- **Some parameters are fixed priors**. Learning interaction strengths from data could improve accuracy but requires sufficient samples per pathway.
- **Temporal dynamics are ignored**. Dynamic networks could model time-series expression data.

PathWeigh II maintains the practical advantages of PathWeigh—open source, extensible Python implementation, support for custom pathways—while providing rigorous treatment of pathway structure. The modest computational overhead (~1.5× runtime) is justified by improved biological validity, particularly for pathways with feedback regulation.

## 5. Conclusion

PathWeigh II demonstrates that probabilistic network inference provides a principled framework for pathway activity analysis. By treating pathways as probabilistic graphical models and employing belief propagation, we overcome limitations of previous methods while maintaining computational efficiency. The integration of gene-level evidence throughout the network, rather than only at inputs, better captures pathway perturbations. We anticipate PathWeigh II will be valuable for systems biology applications requiring robust pathway characterization, particularly in cancer genomics and drug response prediction where regulatory feedback is prevalent.

## Availability

**Source code**: https://github.com/zurkin1/PathWeigh/tree/master/v2 (MIT License)

**Requirements**: Python 3.8+, pandas, numpy, scipy, scikit-learn, networkx

**Data**: Test datasets and pathway database included in repository

## Acknowledgments

This work builds upon research conducted during the author’s doctoral studies at The Mina & Everard Goodman Faculty of Life Sciences, Bar-Ilan University.

